# High-resolution structures of malaria parasite actomyosin and actin filaments

**DOI:** 10.1101/2020.07.02.183871

**Authors:** Juha Vahokoski, Lesley J. Calder, Andrea J. Lopez, Justin E. Molloy, Peter B. Rosenthal, Inari Kursula

## Abstract

Malaria is responsible for half a million deaths annually and poses a huge economic burden on the developing world. The mosquito-borne parasites (*Plasmodium* spp.) that cause the disease depend upon an unconventional actomyosin motor for both gliding motility and host cell invasion. The motor system, often referred to as the glideosome complex, remains to be understood in molecular terms and is an attractive target for new drugs that might block the infection pathway. Here, we present the first high-resolution structure of the actomyosin motor complex from *Plasmodium falciparum*. Our structure includes the malaria parasite actin filament (*Pf*Act1) complexed with the myosin motor (*Pf*MyoA) and its two associated light-chains. The high-resolution core structure reveals the *Pf*Act1:*Pf*MyoA interface in atomic detail, while at lower-resolution, we visualize the *Pf*MyoA light-chain binding region, including the essential light chain (*Pf*ELC) and the myosin tail interacting protein (*Pf*MTIP). Finally, we report a bare *Pf*Act1 filament structure at an improved resolution, which gives new information about the nucleotide-binding site, including the orientation of the ATP/ADP sensor, Ser15, and the presence of a channel, which we propose as a possible phosphate exit path after ATP hydrolysis.

**Significance statement:** We present the first structure of the malaria parasite motor complex; actin 1 (*Pf*Act1) and myosin A (*Pf*MyoA) with its two light chains. We also report a high-resolution structure of filamentous *Pf*Act1 that reveals new atomic details of the ATPase site, including a channel, which may provide an exit route for phosphate and explain why phosphate release is faster in *Pf*Act1 compared to canonical actins. *Pf*Act1 goes through no conformational changes upon *Pf*MyoA binding. Our *Pf*MyoA structure also superimposes with a recent crystal structure of *Pf*MyoA alone, though there are small but important conformational changes at the interface. Our structures serve as an excellent starting point for drug design against malaria, which is one of the most devastating infectious diseases.

## Introduction

Malaria parasites belong to the phylum *Apicomplexa*, which are all obligate intracellular parasites many of which infect humans and livestock, causing immeasurable human tragedy and vast economic losses worldwide. Among the best-known members of the phylum are *Plasmodium* spp. (the causative agents of malaria), *Toxoplasma gondii* (toxoplasmosis), and *Cryptosporidium* spp. (gastrointestinal and respiratory cryptosporidiosis), all life-threatening human pathogens. Apicomplexan parasites are critically dependent on an actomyosin based motor system that drives gliding motility that they use to enter and exit cells of the host organisms (1, 2). The motor complex, called the glideosome, is located in a narrow space between the parasite plasma membrane and an inner membrane complex (IMC), which is structurally unique to this class of parasites (1-3).

Myosins are a large and diverse family of motor proteins, with >30 classes, found across all eukaryotic organisms (4). Although the myosin classes are functionally and structurally distinct, they share several conserved domains (5, 6) and produce force and movement *via* the same basic mechanism, making cyclical interactions with actin, coupled to the break-down of ATP to ADP and inorganic phosphate (P_i_). Most myosins characterized to date move towards the plus-end of actin filaments, with the exception of the reverse-directed, class VI myosins. The myosin heavy chain consists of an N-terminal motor domain that is well-conserved and contains the catalytic site and actin binding site, which attaches and releases actin during the ATPase-cycle. The motor domain is followed by a “neck” region, which binds one or more calmodulin-like light chains. This region is functionally important because structural rearrangements within the motor domain cause it to rotate through a large angle (>60°), and it acts as a lever arm to amplify the motion, producing the active “power stroke”. Class VI myosins have a unique insert within the neck sequence that reverses the power stroke direction. Following the neck region, most myosins have an extended, C-terminal “tail”, which is highly diverse in both sequence and structure and is responsible for targeting the motor to its cellular cargo. Some myosins have a coiled-coil forming region within the tail that causes the heavy chains to dimerize (7).

*Plasmodium* spp. have six myosin genes; three parasite-specific class XIV myosins, two reverse-directed class VI myosins, and one class XXII myosin (8). One of the class XIV myosins, *Pf*MyoA, is essential for gliding motility and is the best-characterized of the parasite myosins. It is unusually small, consisting of just the canonical motor domain and neck region that bears two light chains; essential light chain (*Pf*ELC) and myosin tail interacting protein (*Pf*MTIP) (9). As *Pf*MyoA completely lacks a tail region, it is thought to be unable to dimerize on its own. *Pf*MyoA is anchored to the IMC by binding two glideosome-associated proteins (*Pf*GAP45 and *Pf*GAP50) *via* MTIP (10). To generate gliding motility, *Pf*MyoA interacts with *Pf*Act1 filaments (11-13) that are linked to the plasma membrane *via* the glideosome-associated connector (GAC; (14)), which binds to transmembrane adhesins of the thrombospondin-related anonymous protein family (15). During gliding, *Pf*MyoA moves along actin, pulling the IMC towards the anterior (front end) of the parasite, while pushing the actin filaments and associated plasma membrane, rearwards (16). Recently, X-ray crystal structures have been determined for the truncated motor domains of two *Pf*MyoA orthologs from *T. gondii* (*Tg*MyoA) and *P. falciparum* (*Pf*MyoA) (17, 18). For brevity, herein we refer to the *P. falciparum* proteins without the species-specific prefix “*Pf*”.

Apicomplexan actins differ from the well-characterized canonical actins from yeasts, plants, and animals in some important aspects. While canonical actins form filaments, which can be micrometers to tens of micrometers in length, apicomplexan actin filaments tend to be short, typically around 100 nm (13). For *T. gondii* actin, an isodesmic polymerization mechanism was suggested to be the reason behind the short filament length (19). However, we have shown that the polymerization pathway and kinetics of *P. falciparum* actin 1 (Act1) are very similar to canonical actins (20), and the short length of the filaments is likely caused by a high fragmentation rate (21). In a recent study based on TIRF microscopy, it was suggested that the critical concentration for polymerization of Act1 is at least an order of magnitude higher than that of canonical actins (22). Also the link between ATP hydrolysis and polymerization seems to be different in apicomplexan compared to canonical actins (11, 20, 21). Whereas canonical actins polymerize preferably in the ATP form and only catalyze ATP hydrolysis in the filamentous (F) form, ATP hydrolysis in the monomer causes *Plasmodium* actins to form short oligomers (11). Furthermore, Act1 forms stable dimers, which does not occur in other actins characterized to date (20). These differences indicate that there may be differences in the catalytic mechanism of ATP hydrolysis and phosphate release and in the activation of polymerization between the apicomplexan and higher eukaryotic actins.

Malaria parasites go through several transformations between very different cell types during their life cycle, and some of these forms are among the fastest moving cells. Merozoites penetrate red blood cells in seconds, the midgut-penetrating ookinetes can move at ∼5 µm/min (23) and the mosquito-transmitted sporozoites move with impressive average speeds of 1-2 µm/s (Hopp *et al.* 2015) for tens of minutes. Recently, it was proposed that the speed and force generation of MyoA are fine-tuned by phosphorylation of Ser19 in the N-terminal helix (18). In addition, the light chains are required for maximal speed as measured by *in vitro* actin-gliding assays (9, 24). Structures of the full-length MyoA:ELC:MTIP complex are necessary for understanding the role of the light chains in determining force, speed, and step size of the motor complex as well as its association with other glideosome components.

Here, we report the first structure of full-length MyoA, with its native ELC and MTIP light chains, bound to the Act1 filament and a new high-resolution structure of the bare Act1 filament. Our high-resolution structures show the binding mode of MyoA to Act1, the positioning of the two light chains (ELC and MTIP) on the short MyoA neck, and also reveal a possible route for the leaving phosphate after ATP hydrolysis in the Act1 active site.

## Results and discussion

### High-resolution structures of malaria parasite actomyosin and actin filaments

We reconstituted the malaria parasite actomyosin filament in the rigor state *in vitro* using recombinantly expressed Act1 and MyoA:ELC:MTIP complex in the presence of a small cyclic peptide, jasplakinolide, which stabilizes actin filaments and has been used for cryo-EM studies on undecorated Act1 filaments previously (11, 12). After plunge-freezing and cryo-EM imaging, we observed three types of filaments: bare Act1 filaments and filaments either partially or fully decorated by the MyoA:ELC:MTIP complex. By picking well-decorated filaments from the micrographs **(Fig. S1)**, we determined the malaria parasite actomyosin structure using helical averaging methods (25) to an overall resolution of 3.1 Å, as estimated by Fourier shell correlation with the 0.143 criterion **(Figs. 1 and S1, Table 1)**. For direct comparison, we also determined the structure of the bare Act1 filament at an average resolution of 2.6 Å **(Figs. 1 and S2, Table 1)**. Visual inspection of the density maps of both complexes agrees with the global resolution estimations **(Figs. 1, S3, and S4 and Movies S1-S4)**. We modelled a filament consisting of five Act1 and four MyoA (in the actomyosin filament) subunits modelled into the density **(Fig. S5)**.

**Table 1.** Data collection and refinement statistics.

|  |  |  |  |
| --- | --- | --- | --- |
| 608 | **Data collection** | **Act1-MyoA** | **Act1** |
| 609 | Magnification | 75000 x | 75000 x |
| 610 | Defocus range ($\mu\text{m}$) | 0.8-2.6 | 0.8-2.6 |
| 611 | Voltage (kV) | 300 | 300 |
| 612 | Microscope | Titan Krios | Titan Krios |
| 613 | Detector | Falcon 3 | Falcon 3 |
| 614 | No. of frames | 46 | 46 |
| 615 | Pixel size ($\text{\AA}/\text{pixel}$) | 1.09 | 1.09 |
| 616 | Electron dose ($\text{e}^-/\text{\AA}^2$) | 49.22 | 52.4 |
| 617 | No. of micrographs | 2786 | 3953 |
| 618 | **Reconstruction (Relion)** |  |  |
| 619 | Software | Relion 2.1/3beta | Relion 3.0.7 |
| 620 | Segments | 239225 | 305480 |
| 621 | Box size (px) | 512 | 328 |
| 622 | Rise ($\text{\AA}$) | 28.34 | 28.37 |
| 623 | Azimuthal rotation ($^\circ$) | -168.48 | -167.65 |
| 624 | Average resolution ($\text{\AA}$) (FSC=0.143) | 3.1 | 2.57 |
| 625 | Model resolution ($\text{\AA}$) (FSC=0.5) | 3.2 | 2.7 |
| 626 | Map sharpening B-factor ($\text{\AA}^2$) | 104 | 67 |
| 627 | **Model building (Phenix)** |  |  |
| 628 | No. of atoms | 24052 | 15690 |
| 629 | No. of amino acid residues | 3020 | 1860 |
| 630 | No. of water atoms |  | 140 |
| 631 | No. of ligand atoms ($\text{Mg}^{2+}$, ADP, JAS) | 12 | 15 |
| 632 | Average B-factor ($\text{\AA}^2$) | 62 | 37 |
| 633 | Average B-factor for ligand atoms ($\text{\AA}^2$) | 42 | 36 |
| 634 | Average B-factor for water ($\text{\AA}^2$) | | 31 |
| 635 | R.m.s.d. bond lengths ($\text{\AA}$) | 0 | 0.003 |
| 636 | R.m.s.d. bond angles ($^\circ$) | 0.587 | 0.661 |
| 637 | CC volume | 0.84 | 0.83 |
| 638 | CC masked | 0.88 | 0.87 |
| 639 | Molprobity score | 1.60 | 1.52 |
| 640 | Clash score | 6.08 | 3.51 |
| 641 | Ramachandran plot favoured/allowed/outliers (%) | 96.7/3.3/0 | 98.5/1.5/0 |
| 642 | **Deposition codes** |  |  |
| 643 | PDB | 6TU7 | 6TU4 |
| 644 | EMDB | EMDB-10590 | EMDB-10587 |
| 645 |  |  |  |

**Figure 1.**
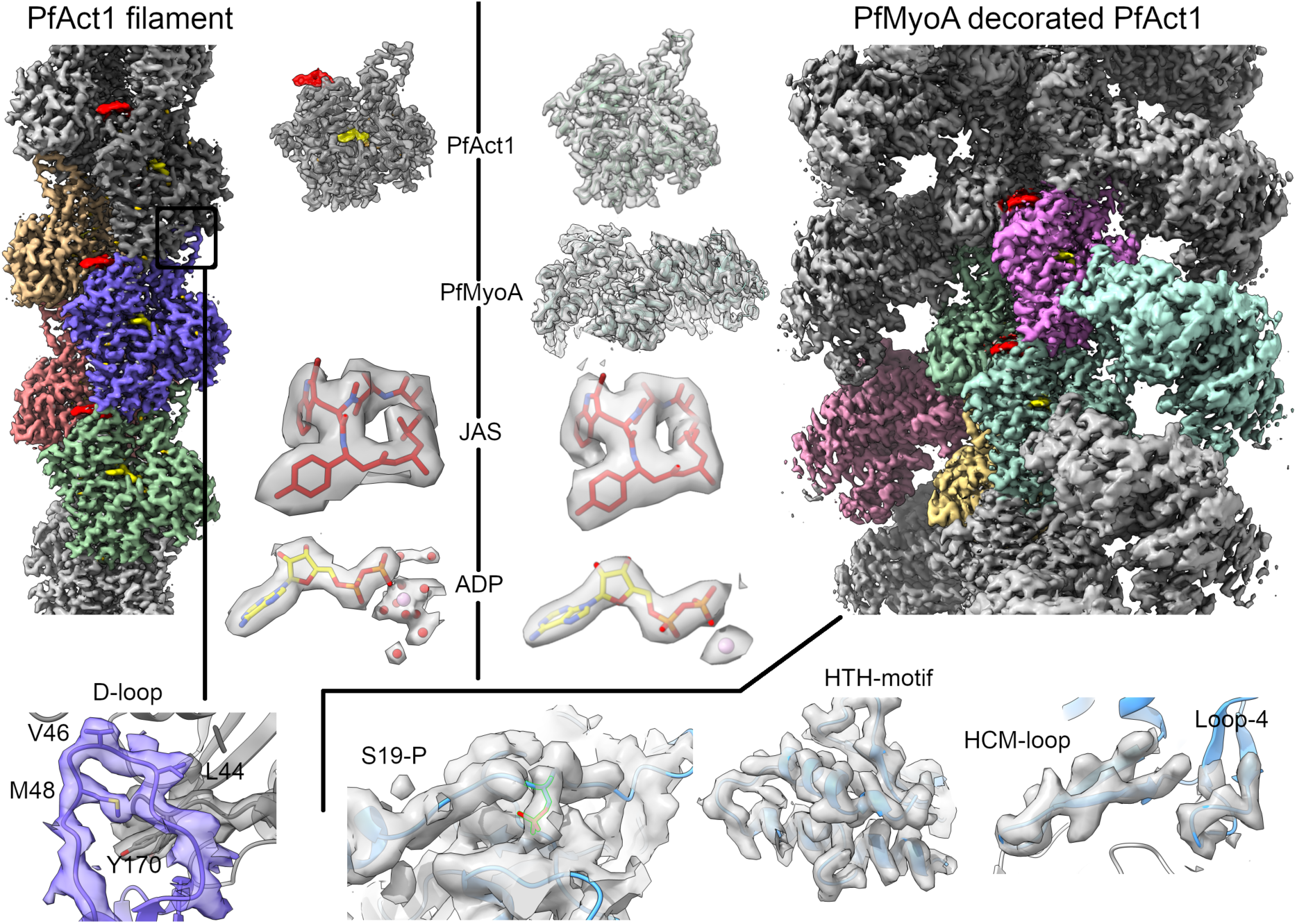
High-resolution reconstructions of filamentous Act1 (left) and MyoA-decorated Act1 (right). In the density maps, the central individual actin protomers and myosins are depicted in different colors. Jasplakinolide (JAS) and ADP are shown in red and yellow, respectively. The top middle part shows density maps around individual protein molecules and small molecule ligands. The bottom row shows density maps of selected motifs, as labeled, within the interfaces in the complexes. Additionally, density around phosphorylated Ser19 (S19-P) is shown.

As typical for actomyosin cryo-EM structures (26-29), the resolution in the Act1:MyoA:ELC:MTIP structure decays radially away from the helical axis, having a well-resolved core, consisting of the actin filament and the myosin motor domain, whereas density for the MyoA neck region is weaker, but nevertheless allowed us to locate density for the light chains, ELC and MTIP (**Fig. 2**). The actin core in both of our structures is particularly well resolved compared to other actin filament structures **(Fig. S3, S4)**. Thus, the molecular details of F-actin, including the small molecules ADP and jasplakinolide as well as the active site metal and several water molecules, can be visualized in greater detail **(Fig. 1)** than in myosin-decorated or undecorated actin filament structures to date (12, 26-31). The majority of the MyoA motor domain is well-ordered, and most of the side chains can be placed into density, especially within the actomyosin interface, allowing us to analyze specific contacts between MyoA and Act1 in detail.

**Figure 2.**
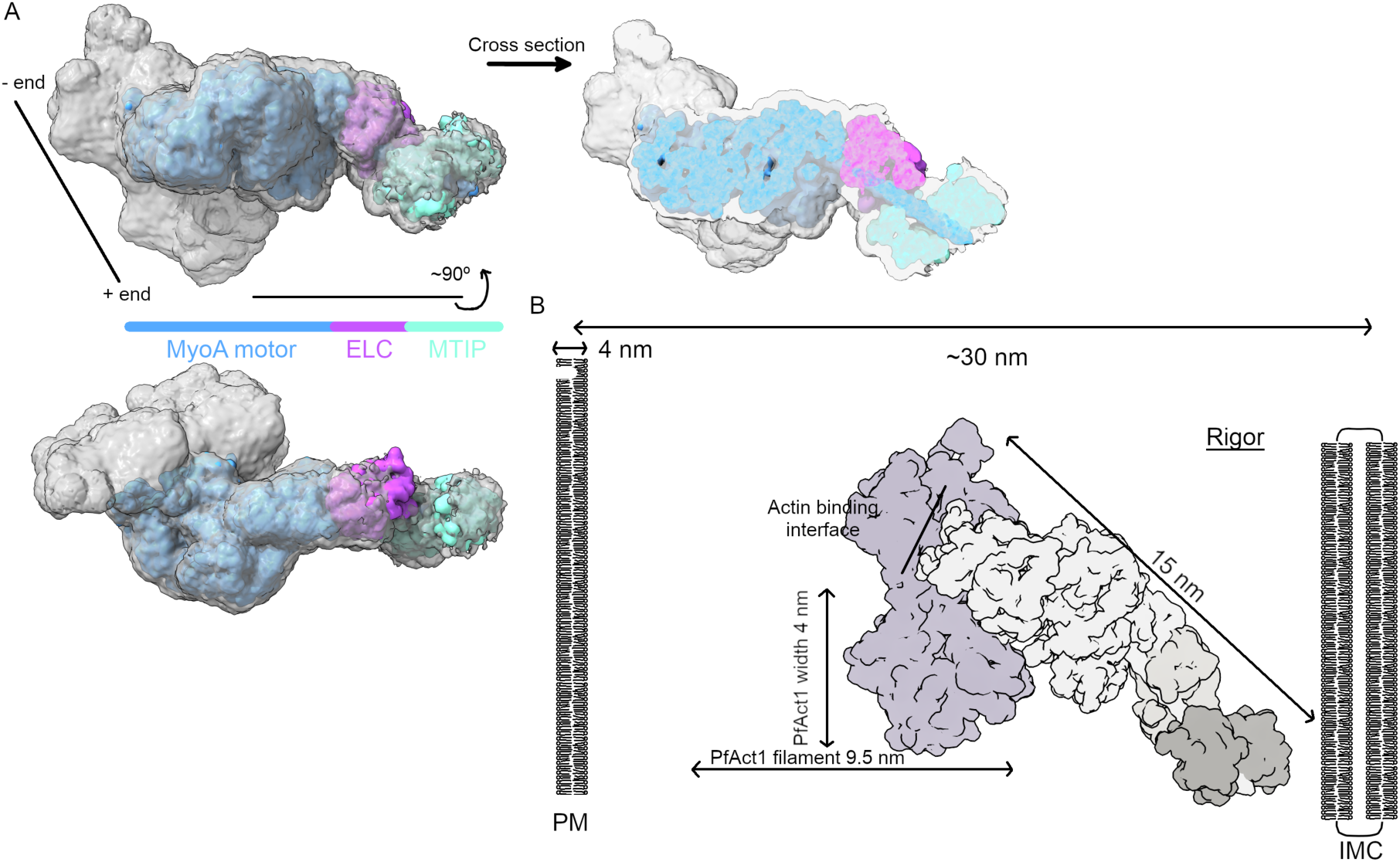
A model of the light chains ELC and MTIP bound to MyoA. (A) An unsharpened map (gray) shows the location of the MyoA motor (blue) and the light chains ELC (magenta) and MTIP (cyan). (B) The Act1:MyoA:ELC:MTIP model in the rigor state shown in the context of the membrane-delimited sub-pellicular compartment.

ELC and MTIP bind to the short neck of MyoA in a compact manner, presumably stiffening the neck region so that it acts as a relatively rigid lever arm **(Fig. 2)**, consistent with the requirement of both light chains for maximal actin gliding speed as measured in *in vitro* motility assays. A consequence of the rigid neck region is that the space taken up by the actomyosin complex in the sub-pellicular compartment must remain rather fixed. The length of the MyoA:ELC:MTIP complex would occupy approximately half the distance between the IMC and the plasma membrane (∼30 nm; **Fig. 2**), which is similar to the spacing between thick and thin filaments in the muscle sarcomere (32). In the parasite, when the additional space taken up by the actin filament (diameter ∼6.5 nm) and GAC (D_max_ ∼25 nm; (14)) is considered, there are significant geometric and steric constraints on how the actomyosin complex might be arranged. It is possible that the attachment of MyoA *via* its MTIP light chain to the GAP45:GAP50 complex in the IMC might allow azimuthal movement so the motor domain might approach the actin helix from different angles. These structural considerations will need to be revisited when the 3D geometry of the full glideosome complex is understood.

The MyoA motor domain has a conserved overall structure with several known subdomains common to functional myosin motors **(Fig. S6)**. The upper and lower 50 kDa subdomains are separated by the actin-binding cleft, which is in a closed conformation in our complex **(Fig. 3)**. The central transducer region consists of seven β strands **(Fig. S6)**. The lower 50 kDa subdomain contains a helix-turn-helix (HTH) motif, involved in intimate contacts with Act1, while the rest of the actin-contacting residues are in the hypertrophic cardiomyopathy (HCM) loop and in loops 2, 3, and 4 **(Fig. 3)**. The more distal parts, the converter and the relay helix, are involved directly in the conformational change upon the power stroke **(Fig. 2, (33))**. Additionally, class XIV myosins have an N-terminal SH3 domain, the role of which is unclear but could be related to the regulation of myosin function during different stages of the parasite life cycle (34). Our actomyosin rigor structure shows that neither the SH3 domain nor the N-terminal helix is in direct contact with the actin filament **(Fig. 3)**.

**Figure 3.**
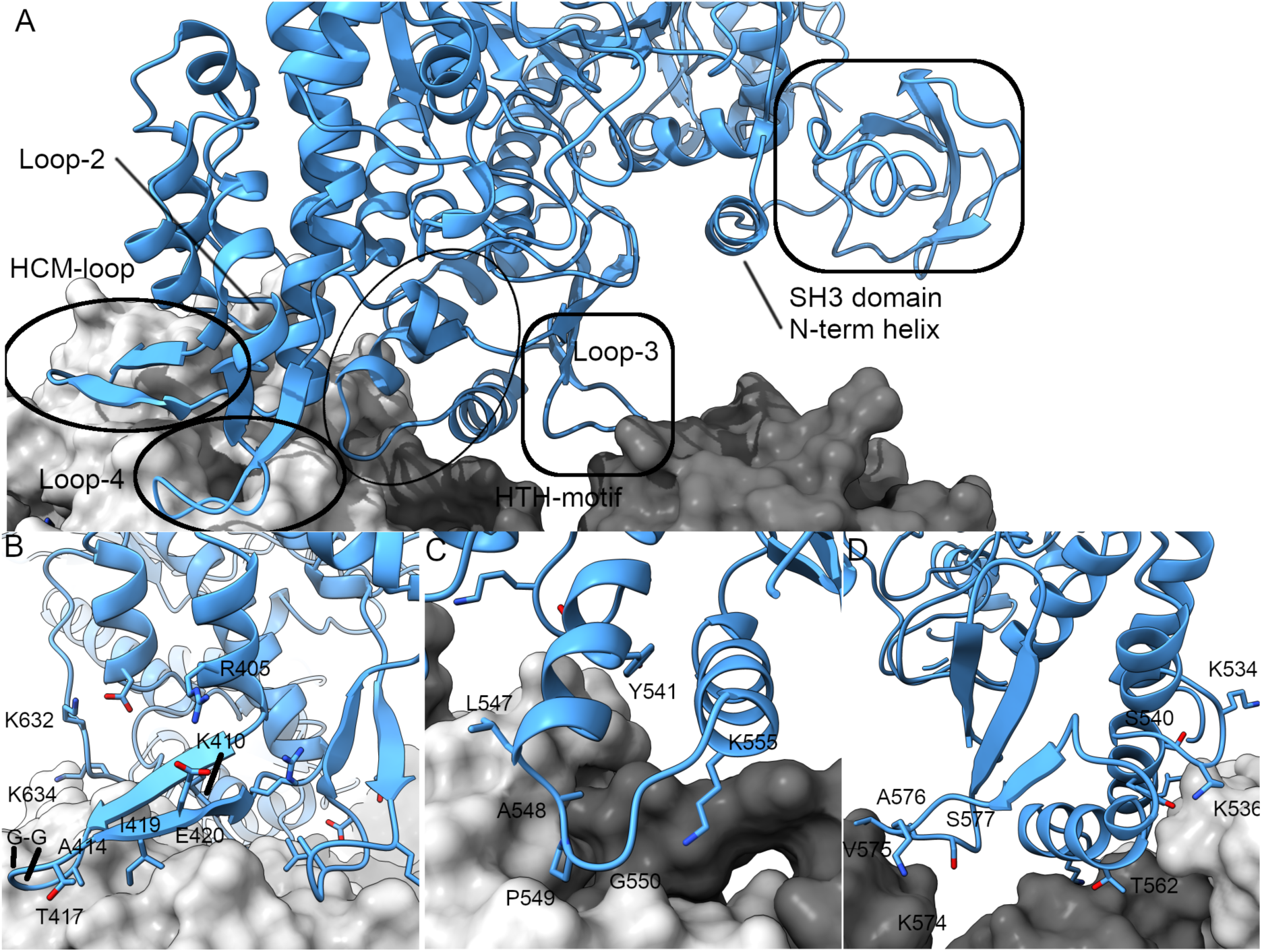
Molecular interactions of MyoA with Act1. (A) MyoA is shown as blue cartoon and ActI as gray surface representation. The two adjacent actin protomers are colored with different shades of gray. The MyoA motifs contributing to the interface as well as the N-terminal helix and SH3 domain are highlighted and labeled. (B) The HCM-loop lies on one actin protomer, contacting SD1. The basic loop 2 (Lys632 and Lys634) interacts with the acidic N-terminus of Act1 (C) The HTH motif in the lower 50K domain contributes the majority of specific hydrogen bonds in the interface, interacting with the D-loop (SD2) and the C-terminal patch of Act1 between SD1 and SD3 in close vicinity of the D-loop in the adjacent actin subunit. (D) Loop 3 and the HTH motif visualized approximately 180° rotated with respect to panel C. Loop 3 (576-577) interacts with the adjacent actin monomer on SD1. The activation loop in the lower 50K domain protrudes towards the next SD1, Lys534 interacting with actin N-terminus.

Overall, the MyoA structure in complex with actin is very similar (with an overall root mean square deviation (r.m.s.d.) for the C_α_ positions ∼1.3 Å) to the rigor-like conformer found as one of the molecules in the asymmetric unit in the recently determined MyoA crystal structure (18). However, there are several small but important changes in the complex. Notable differences allow interactions and avoid clashes with actin, including movement of the HCM loop (Ile409-Arg422) by nearly 4 Å, slightly smaller shifts in loop 3 (Glu369-Ala378), loop 4 (Thr571-Lys579), and the HTH motif (Val538-Asn563) as well as the ordering of loop 2 (Lys632-Ser639), which is not visible in the crystal structure (18). In addition, there are several smaller rearrangements in more distal areas, in particular in the converter domain. The N-terminal helix, which is specific to apicomplexan MyoAs and accommodates the phosphorylated Ser19, is stacked between the SH3 domain and the motor domain, suggesting an intimate structural or functional role **(Figs. 3 and S6)**. In the crystal structure of TgMyoA, the entire N-terminal helical extension is not visible in the electron density map (35) but is seen in the most recent MyoA crystal structures (18).

### Actomyosin interface

The MyoA domain architecture is conserved, as in other myosins, regardless of the myosin class **(Figs. 2 and 3)**, but residues in the actin-binding interface are generally not conserved (**Fig. S6)**. The Act1 surface is relatively flat, and thus MyoA also lacks large protrusions, which would penetrate deep into the filament **(Fig. 3A)**. The interface is mainly formed by contacts between the HTH motif, the HCM loop, and loops 2, 3, and 4 **(Fig. 3)** similarly to previous structures (28, 29).

The majority of specific contacts are formed by the HTH motif in the lower 50K domain, which contacts the actin-actin interface (buried surface area 386 Å^2^), where the D-loop of one subunit inserts into the cleft between subdomains 1 and 3 of an adjacent subunit **(Fig. 3B and C)**. The first half of the HTH motif (Ser540-Gly550) forms extensive contacts with both actin subunits, whereas the latter part (Gly551-Lys565) interacts mostly with the D-loop. The Asp544 side chain of MyoA forms hydrogen bonds with the backbone amine of Thr352 in Act1 and the side chain or backbone amine of Ser351. Asp544 also interacts *via* its backbone carbonyl with the backbone amine of Ala548 in MyoA. This interaction network between Act1 and MyoA occurs just before the kink in the relay helix, possibly stabilizing it. The tip of the HTH motif (Ala548-Pro549-Gly550) inserts into a hydrophobic cleft in Act1, lined by Ile347-Leu352 and Tyr145-Thr150 as well as the D-loop residues Met46-Val47 from the adjacent actin subunit.

The HCM loop **(Fig. 3B)** forms a second important interface to Act1 (buried surface area 295 Å^2^) that is complementary in shape and hydrophobic in nature. Suprisingly, there are no side chain interactions contributing to this interface, which is rather mediated by main chain atoms. Additional contact points are created by loop 2, which is in close vicinity of the HCM loop. The backbone atoms of Gly24 and Asp26 in Act1 form hydrogen bonds with the backbone atoms of Ile635 and Gly633, respectively, in MyoA. Ile635 of MyoA inserts into a small patch on Act1 lined by Gly25, Asp26, Ser345, Ile346, and Ser349. The MyoA basic residues Arg606, Lys634, and Lys637 are close to Asp26, Asp25, and Ser349/Ser351, respectively, distances ranging between 3.8 and 4.5 Å.

### Details of the actin 1 filament and implications on phosphate release

We reconstructed the Act1 filament alone and obtained helical parameters that were highly similar to the Act1:MyoA complex. As previously noted for other actomyosin complexes (26, 27, 29), neither MyoA nor Act1 undergo large conformational changes when compared to the structures in isolation. However, our highly-resolved structures of the Act1 filament allow us to see most of the side chain orientations and details of the binding of the ligands, ADP-Mg and jasplakinolide, as well as putative water molecules, including several in the vicinity of the active site **(Fig. 1, Movies S3 and S4)**.

Based on crystal structures, the side chain of Ser15 (14 in canonical actins) has been established as a nucleotide sensor, which is turned away from bound ATP but hydrogen bonded to the β-phosphate of ADP (36). In our structure of the ADP-bound form, the side chain of Ser15 points away from the nucleotide, not contacting the ADP phosphates, thus more closely resembling the conformation in crystal structures of the ATP state **(Fig. 4)**. Interestingly, the Ser15 side chain points towards a channel with additional density, which in our high-resolution maps is close to where P_i_ is bound in the recent F-actin structures in the ADP-P_i_ state (37). The channel extends towards solvent, but its opening is partly blocked by the side chains of Asn116 and Trp80 **(Fig. 4A and B, Movie S4)**. We have modelled this density as a chain of water molecules **(Figs. 4 and S7)**, but it cannot be excluded that the channel could be partly occupied by for example phosphate or other solute molecules. The corresponding channel in monomeric Act1 is much shorter compared to the F-form and does not extend to solvent **(Fig. 4C)**. In F-form, this channel appears to be the most favorable way out from the active site. Thus, we propose it may be an alternative path for the leaving phosphate after hydrolysis, instead of or in addition to the proposed backdoor (37). In this pathway, Asn116 and Trp80, would act as “gatekeepers” that switch conformation to allow the phosphate to leave, as simply changing the rotamers of these side chains is sufficient to fully open the channel to solvent **(Fig. 4D)**.

**Figure 4.**
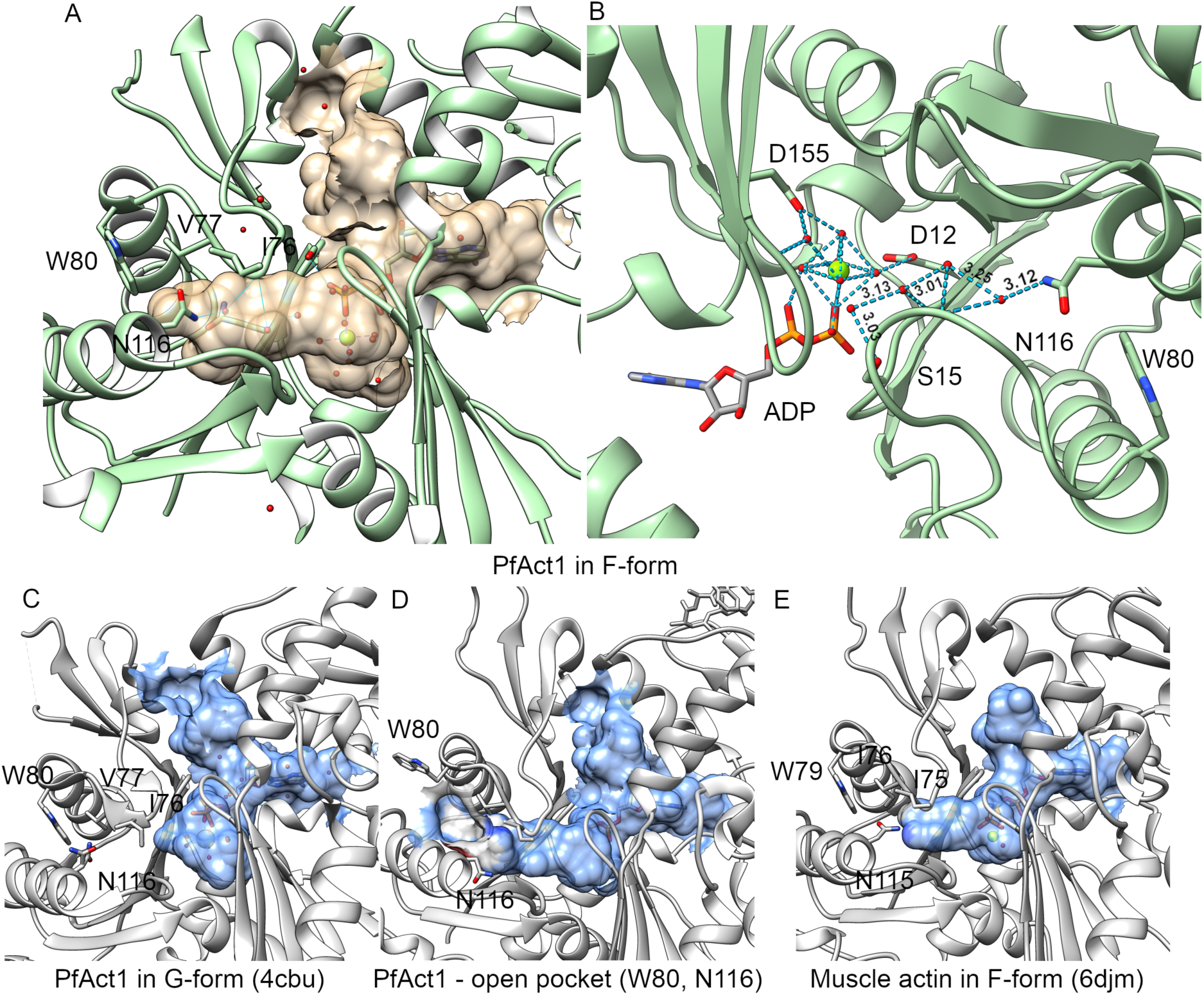
Internal cavities in filamentous and monomeric actin structures. The internal cavities were calculated using the CASTp server (60) and visualized as transparent surfaces using Chimera (50). (A) The internal cavity in the PfAct1 filament and (B) the corresponding cavity rotated about 180° with the water hydrogen bonding network depicted. (C) The corresponding channel in monomeric PfAct1 (4cbu; (11)). (D) The channel in the PfAct1 filament becomes solvent accessible upon rotation of Trp80 and Asn166 side chains to alternative rotamers. (E) The corresponding cavity in muscle actin (6djm; (38)) is shorter due to a conservative amino acid change from Ile76 to V77 in PfAct1.

Similar density as seen in our structure has been described at this site in the recent AMP-PNP and ADP-P_i_ structures of muscle actin (38). In muscle actin, also in F-form, the cavity is, however, much shorter due to the presence of the bulkier Ile76 instead of V77 in Act1 **(Fig. 4E)**; phosphate release would be restricted and could possibly be an additional reason for the differences in phosphate release rates in different actins. Notably, residue 76/77 is an Ile in α- and γ-actins, but is also Val in β-actin as well as plant and yeast actins. We have previously noticed the importance of the substitution of the residue preceding Asn116 from an Ala in canonical actins to Gly in Act1. Ala at this position is responsible for structuring the C terminus and slightly slows down phosphate release compared to a Gly, in particular in the Mg-bound form (11, 21). In addition to allowing more flexibility to the C terminus, a Gly at this position would enable more conformations for Asn116 and thus faster opening of the phosphate release channel, contributing together with the movement of the A-loop (21) to the faster P_i_ release in the parasite actins. In concordance with this, jasplakinolide, which inhibits phosphate release and stabilizes filaments, is bound very close to this proposed exit route.

Interestingly, Asn115 and Arg116 in skeletal or smooth muscle α-actin are sites of disease-associated mutations (N115S, N115T, R116Q, R116H). The mechanisms by which these mutations disrupt actin function are not known but have been postulated to be linked to nucleotide binding or association with actin-binding proteins, such as CAP or α-actinin (39, 40). The mutations N115T and R116Q in yeast actin affect both nucleotide exchange as well as polymerization kinetics, in particular in the nucleation phase (41).

## Concluding remarks

The first malaria parasite actomyosin filament structure reported here will serve as a foundation for understanding gliding motility in Apicomplexa as well as force generation in a variety of actomyosin motor systems. We visualize the actin-myosin interface at high-resolution and also present the structure of the entire core motor complex including the light chains which are essential for fast motility. The structure of the complex will also be an important tool for evaluating the actomyosin motor as a drug target and describes several protein-protein interfaces that may be targeted by inhibitors. The bare Act1 filament assigns accurate side chain conformations and reveals new active site features including a channel, which we propose as a possible release pathway for the leaving phosphate in Act1.

## Materials and methods

### Preparation of proteins and cryo-EM grids

MyoA:ELC:MTIP and Act1 were expressed and purified as previously described (9, 20). For Act1 samples, ammonium acetate was removed by a spin column, and Act1 (13.1 µM) was polymerized in the presence of 13 µM jasplakinolide by adding 10X KMEI (10 mM Hepes, pH 7.5, 500 mM KCl, 40 mM MgCl2, 10 mM EGTA). Polymerized filamentous Act1 was diluted down to 0.13 µM, and treated with apyrase (77 µg/ml) for 20 min before addition of the MyoA:ELC:MTIP complex in a 1:1 ratio. For Act1-only samples, no apyrase was used. After 30 min incubation, negatively stained or cryo-EM grids were prepared. Quality of the decorated filaments was first checked with negative staining, and after confirming presence of decorated Act1 filaments, frozen-hydrated samples were prepared on Quantifoil 2/2, 200 mesh air-glowed grids using an Mk III Vitrobot (FEI). 3 µl sample was applied on a grid and blotted for 3 s before plunge-freezing in liquid ethane.

### Data collection and image processing

#### Actin 1:myosin A filament

Data consisting of 2815 movies were collected with a Titan Krios electron microscope equipped with a Falcon 3 camera, operated at 300 kV in counting mode. The magnification was 75000x, corresponding to 1.09 Åpixel^−1^, using dose rate 0.539 e/Å^2^s^−1^ with exposure time 92 s for a stack of 46 frames. The dose per frame was 1.07 e/Å^2^, and we used defoci from 0.8 to 2.6 µm. The movies were aligned with MotionCor2 (42) within the Scipion framework (43). Contrast transfer function was estimated by CTFFind 4.1.5 (44), and the resulting 2786 good images were used for further data processing. To confirm data processability, we picked decorated filaments by hand and performed low-resolution reconstructions from 2x binned images from a few hundred particles using symmetry parameters defined for Act1 (12) or refined helical symmetry parameters in Relion 2.1 (25, 45) using a featureless cylinder as a starting reference. The reconstruction converged into a filamentous structure with protruding densities typical for actomyosin structures. Further inspection revealed that homologous myosin motor domains (28) as well as a filamentous Act1 model (PDB code: 5ogw (12)) fit the density well. Decorated filaments were iteratively picked with Relion autopick using reference-free Relion 2D classes as templates with an inter-box distance of two subunits with a rise of 28 Å. The resulting 381256 particles were subjected to reference-free 2D classification in Relion, resulting in 239225 fully decorated actomyosin particles. High-resolution volumes were generated in Relion 2.1 auto-refine using low-resolution models from 2x binned data as reference. A soft mask was applied on a reference model, and reconstruction was continued from the last iteration. When the reconstruction converged, we continued refinement of the contrast transfer function and subsequent reconstructions with Relion 3 beta for two additional rounds. Refined half-maps were sharpened automatically (46), and global resolution was corrected for the effects of a mask (47) using the Relion postprocessing tool. The global resolution reached 3.1 Å based on the 0.143 Fourier shell criterion. The local resolution was calculated with Blocres in Bsoft package version 1.8.6 with a Fourier shell correlation threshold of 0.3 (48, 49).

#### Actin 1 filament

We collected 3953 movies with a Titan Krios electron microscope fitted with Falcon III camera in electron counting mode, operated at 300 kV. We used a nominal magnification of 75000x, corresponding to 1.09 Åpixel^−1^. Each movie was recorded as 46 frames with 1.17 e^−^/Å^2^ dose/frame 1.17 e/Å^2^ using defoci from 0.8 to 2.6 µm. The fractionated movies were aligned globally in Relion 3.0.7. Contrast transfer function was estimated by CtfFind 4.1.10 from dose-weighted micrographs. Particles were picked automatically using four times the rise (28 Å). The resulting 336552 particles were extracted and binned two times at 2.18 Å/pixel in a 328 pixel box before they were subjected to a reference free 2D classification, and resulted in 305480 particles in the final data set. These particles were re-extracted as unbinned, and processed with Relion auto-refine with a reference mask and solvent flattening from a previous reconstruction at nominal resolution 4.4 Å low pass filtered to 20 Å. After polishing and contrast transfer function refinement, the resulting map reached an average resolution of 2.57 Å, based on the 0.143 Fourier shell criterion. Visual inspection of the density map features agreed with the resolution estimate. After reconstruction, the map was masked and sharpened by the Relion postprocessing tool.

### Model Building

#### Actin 1:myosin A filament

Act1 (PDB code 5ogw; (12)) was docked in density using Chimera (50), and we generated four additional symmetry copies. Two copies of MyoA crystal structure in the rigor-state (PDB code 6i7d; (18)) were placed into the density using Chimera. We iteratively built a model using tools in ISOLDE 1.0b (51), Coot (52), and real-space refinement in Phenix 1.16 (53). When the model was complete, model geometry was optimized by Namdinator (54, 55). The local resolution of the reconstruction was estimated by Blocres (47) and Chimera and ChimeraX (50, 56).

In addition, we note that the experimental map is of high quality. Before the MyoA crystal structure became available, we built a complete model of MyoA, *de novo*, based on a homology model of MyoA, the density map, and myosin models 1DFK and 6C1D (28, 57) to guide manual model building. MyoA was correctly built into density, and we have assigned the sequence register correctly, excluding the last 25 amino acids, which were better resolved in the X-ray crystal-structure (PDB code: 6i7d). Additionally, phosphorylation of Ser19 was resolved with the correct sequence register.

#### Actin 1 filament

Our previous model of the Act1 filament (PDB code: 5ogw; (12)) was placed in the density map in Chimera (50), and was manually build using model building tools in ISOLDE 1.0b3.dev6 (51), Coot (52), and real-space refinement in Phenix 1.17 (53). We built five copies of Act1, re-assigning side chain rotamers as required, and waters around Act1 monomers into density. Additionally, we built the active site with the Mg^2^-coordinated waters. Density for the Mg^2+^-water complex at the active site was elongated in the plane of the atoms, and the Mg^2+^-coordinated waters were merged together with the Mg^2+^ ion density. Therefore, we created tighter custom restraints to accommodate ideal coordination around the Mg^2+^ ion.

#### Low-resolution model of light-chain-decorated myosin A

I-TASSER server (58) was used to build a homology model of ELC (PF3D7_1017500). The resulting map and model from the high-resolution refinement (see above) worked as a basis for atomic model building. An alanine helix was placed into density with Coot (52) continuing from the last residue (Lys768) of the high-resolution MyoA model to residue 817. The homology model of ELC and a crystal structure of MTIP (residues 60-204) in complex with a MyoA peptide (2QAC, (59)) were placed into density using Chimera (50). The resulting coordinates were refined using Namdinator (54). For analysis, we generated a density presentation of each chain with Molmap low-pass filtered down to 5Å: The ELC, MTIP, MyoA, and their density fits were further optimized manually.

## Acknowledgements

We thank the Francis Crick Structural Biology Scientific Technology Platform for instrument access, Andrea Nans for help with data cryo-EM data collection, and Phil Walker for help with computing. We thank Donald Benton and Oliver Acton for helpful advice with image processing. I.K. and J.V. were supported by grants from the Academy of Finland, the Sigrid Jusélius Foundation, and the Norwegian Research Council. P.B.R. and J.E.M. are supported by the Francis Crick Institute, which receives its core funding from Cancer Research UK (FC001143, FC001178), the UK Medical Research Council (FC001143, FC001119), and the Wellcome Trust (FC001143, FC001119). The maps and models have been deposited in the Electron Microscopy Data Bank, http://www.ebi.ac.uk/pdbe/emdb/ (accession nos. EMD-10590 and EMD-10587). The atomic models have been deposited in the Protein Data Bank, https://www.ebi.ac.uk/pdbe/ (PDB ID codes 6TU7 and 6TU4).

## Movie captions

**Movie 1. The Act1:MyoA map colored by local resolution.** First, the malaria parasite actomyosin complex is rotated around the helical axis to present the overall quality of the map used to build the actomyosin model. The latter part of the movie shows the maps for a longitudinal Act1 dimer and MyoA separately. The slider on the right indicates the color code for the resolution in different parts of the map from 2.5 (blue) to 4.5 Å (red). The Fourier shell correlation threshold was 0.3.

**Movie 2. Quality of the Act1:MyoA map.** First, the MyoA-decorated filament is rotated around the helical axis, demonstrating high-resolution electron density, in which the majority of side chains are visible, sufficient for *de novo* atomic model building. In the latter part of the movie, each of the central protein subunits (4 actins and 2 myosins) is depicted with a different color. Jasplakinolide, located between actin subunits, is shown in red and ADP in the cleft between the actin subdomains is shown in yellow.

**Movie 3. High-resolution map of the Act1 filament colored by local resolution.** First, the filament is rotated around the helical axis, demonstrating high-resolution features. The latter part of the movie shows a single actin protomer. The slider on the right indicates the color code for the resolution in different parts of the map from 2.1 (magenta) to 2.5 Å (cyan). The Fourier shell correlation threshold was 0.143.

**Movie 4. A cross-section of Act1 showing the internal channel extending from the nucleotide binding pocket towards Asn116 and Trp80 on the surface.** The Act1 model is shown as surface representation, and the plane of view sections through the internal void revealing putative waters (red spheres) and ADP (sticks).

