## Supplemental Figures for "High-resolution structures of malaria parasite actomyosin and actin filaments"

**Supplementary figures and legends**

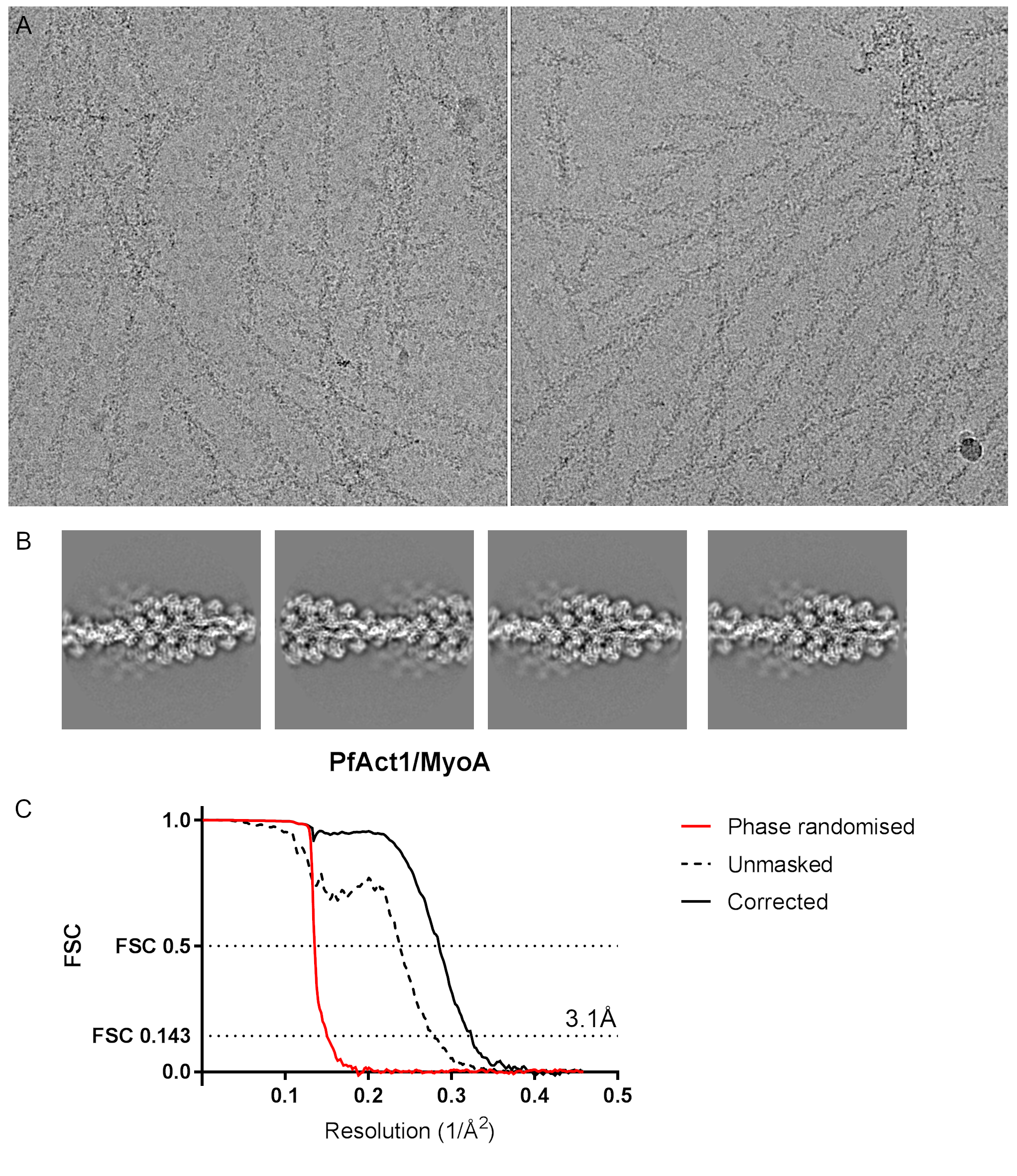

**Figure S1. ActI:MyoA filaments and resolution reconstruction.** (A) Representative micrographs of MyoA-decorated ActI filaments, (B) reference free classes derived from them, and (C) Fourier shell correlation of the ActI:MyoA complex. The masked curve was calculated from independently refined half-datasets with a soft-mask filtered to 15 Å.

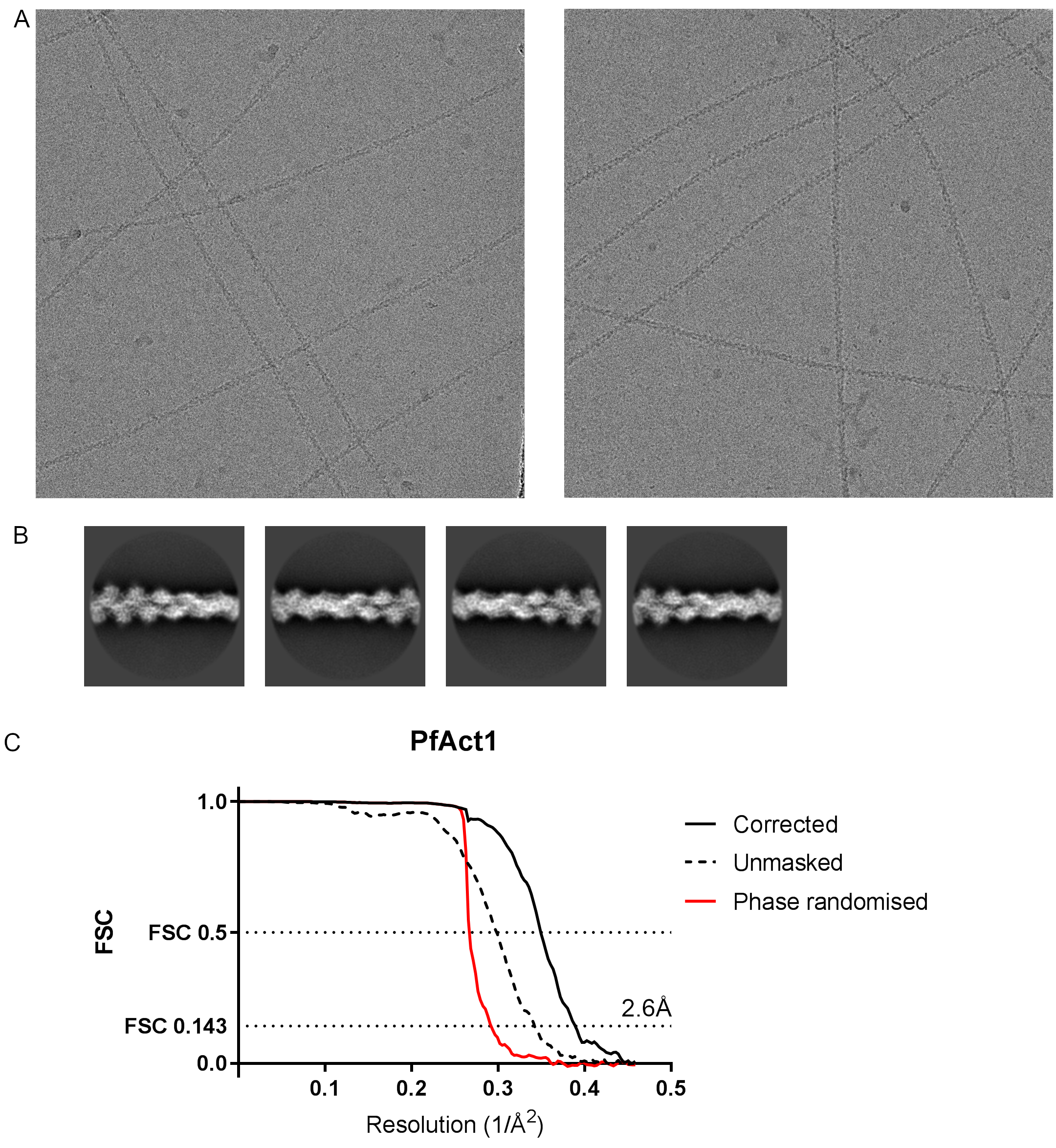

**Figure S2. Examples of the Act1 filaments and resolution reconstruction.** (A) Representative micrographs of ActI filaments low-pass filtered at absolute frequency 0.15, (B) reference free classes derived from them, and (C) Fourier shell correlation of the PfActI filament. The corrected curve was calculated from independently refined half-datasets with a soft-mask filtered to 15 Å in Relion.

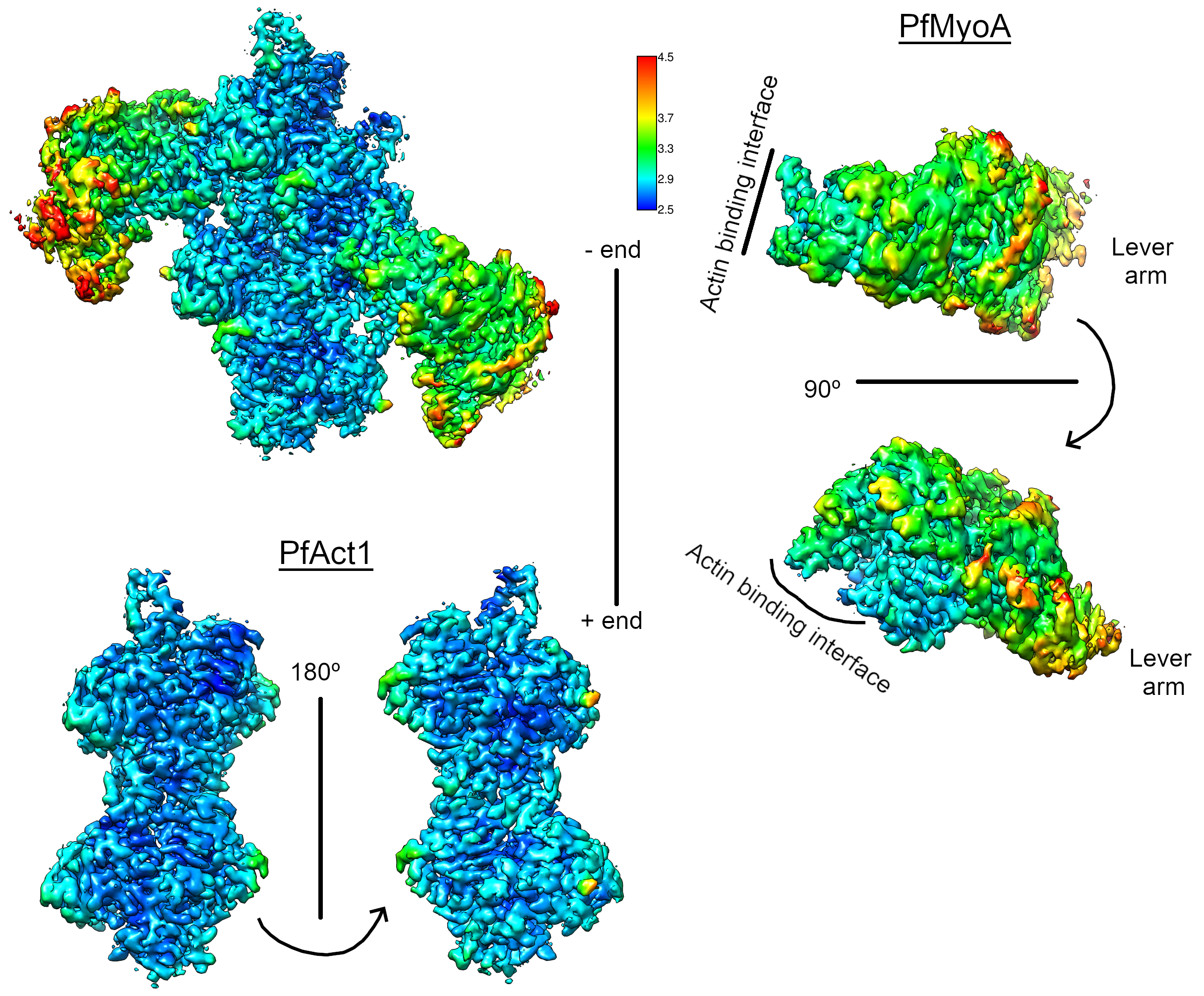

**Figure S3. Local resolution of the MyoA-decorated Act1 filament.** The local resolution estimation is based on the Fourier shell correlation threshold 0.3 calculated with Blocres in the Bsoft software package, which was applied to the final sharpened map.

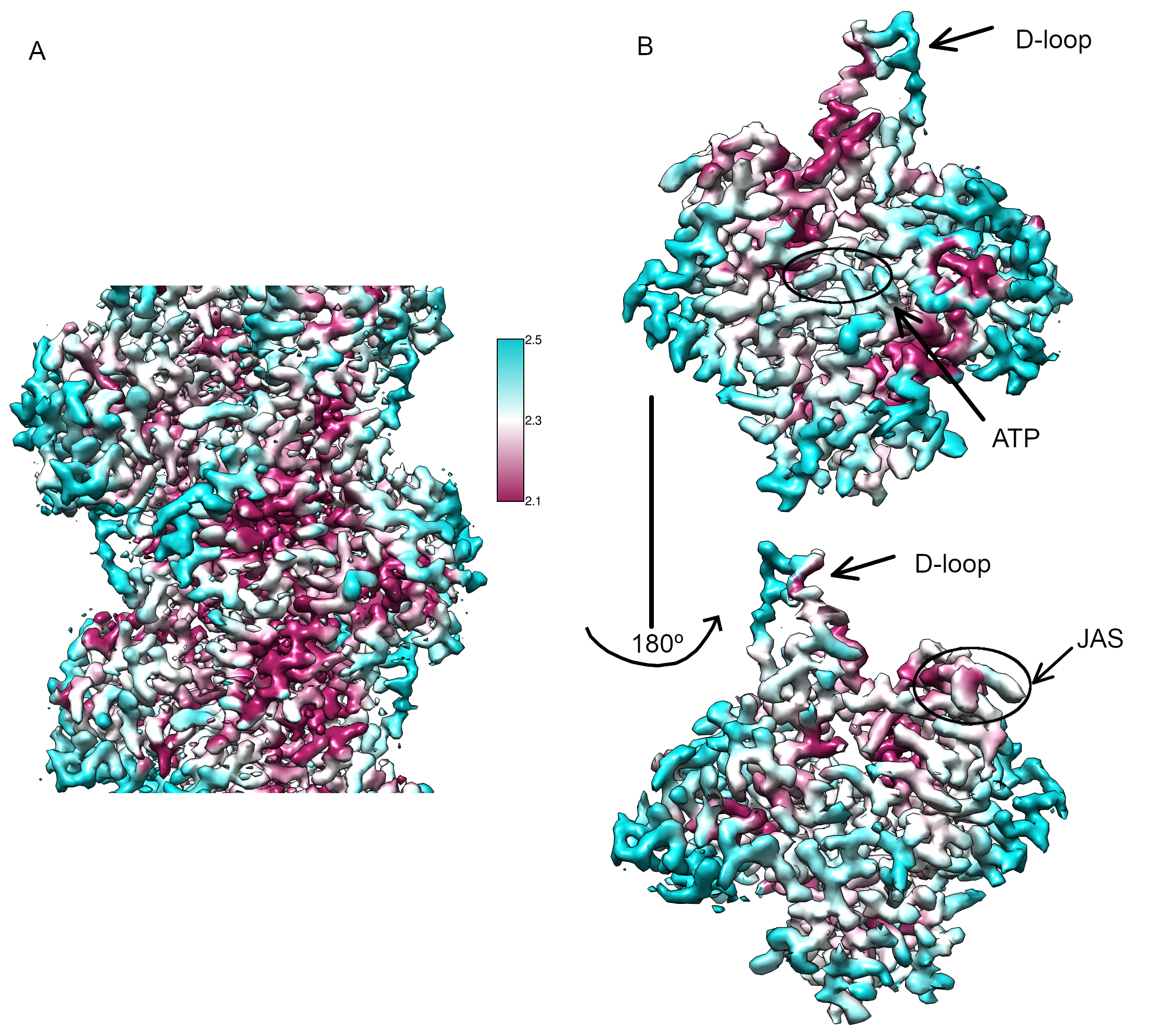

**Figure S4. Local resolution of the Act1 filament.** The local resolution estimation is based on Fourier shell correlation threshold 0.143 calculated with Blocres in the Bsoft software package, applied on the final sharpened map. The left panel shows a central section of the filament, and right panel show an individual actin protomer with the ligand densities highlighted.

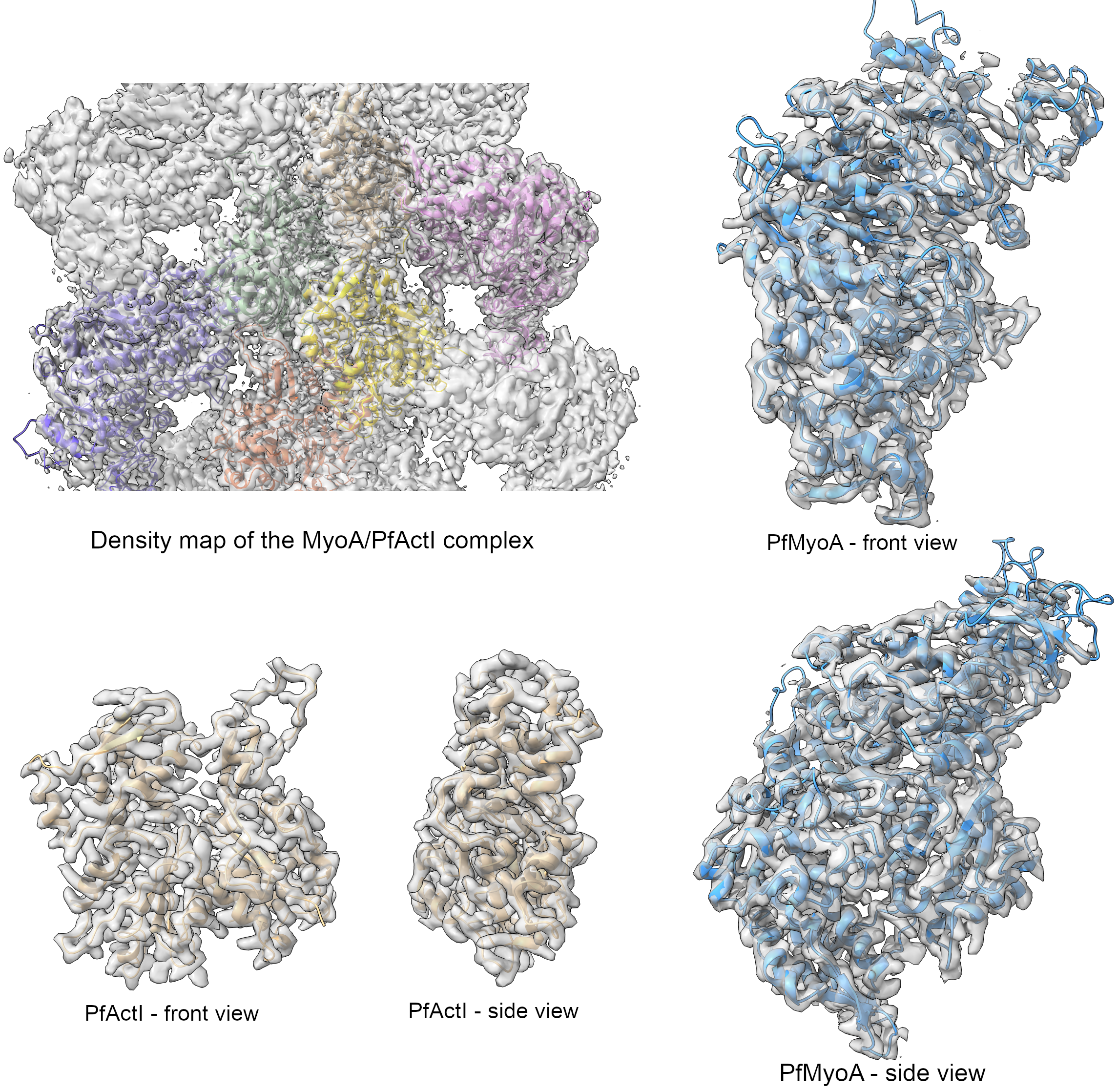

**Figure S5.** Density maps of the Act1:MyoA complex and individual sub-units.

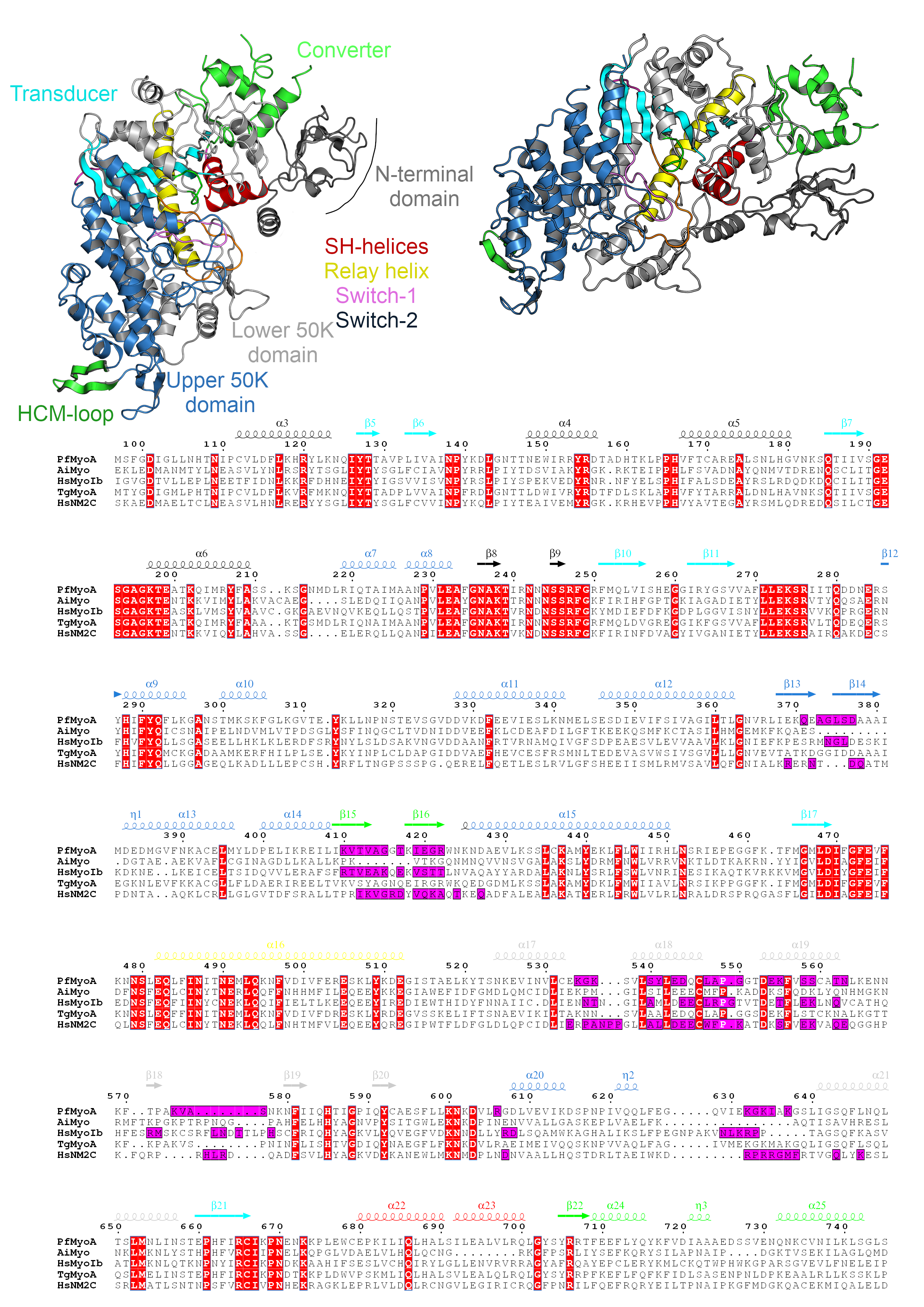

**Figure S6. Structure of the MyoA motor domain.** The MyoA motor domain is depicted as cartoon representation in two different orientations, and the different motifs are shown in different colors and labeled. The coloring of the secondary structure elements above the sequence alignment corresponds to that of the myosin structure above. Strictly conserved residues are boxed with red and interface residues are indicated with purple boxes. The sequence alignment was performed in JalView (61) using default settings and visualized using the ESPript web interface (62). Pf: *Plasmodium falciparum*, Ai: *Argopecten irradians*, Hs: *Homo sapiens*, Tg: *Toxoplasma gondii*.

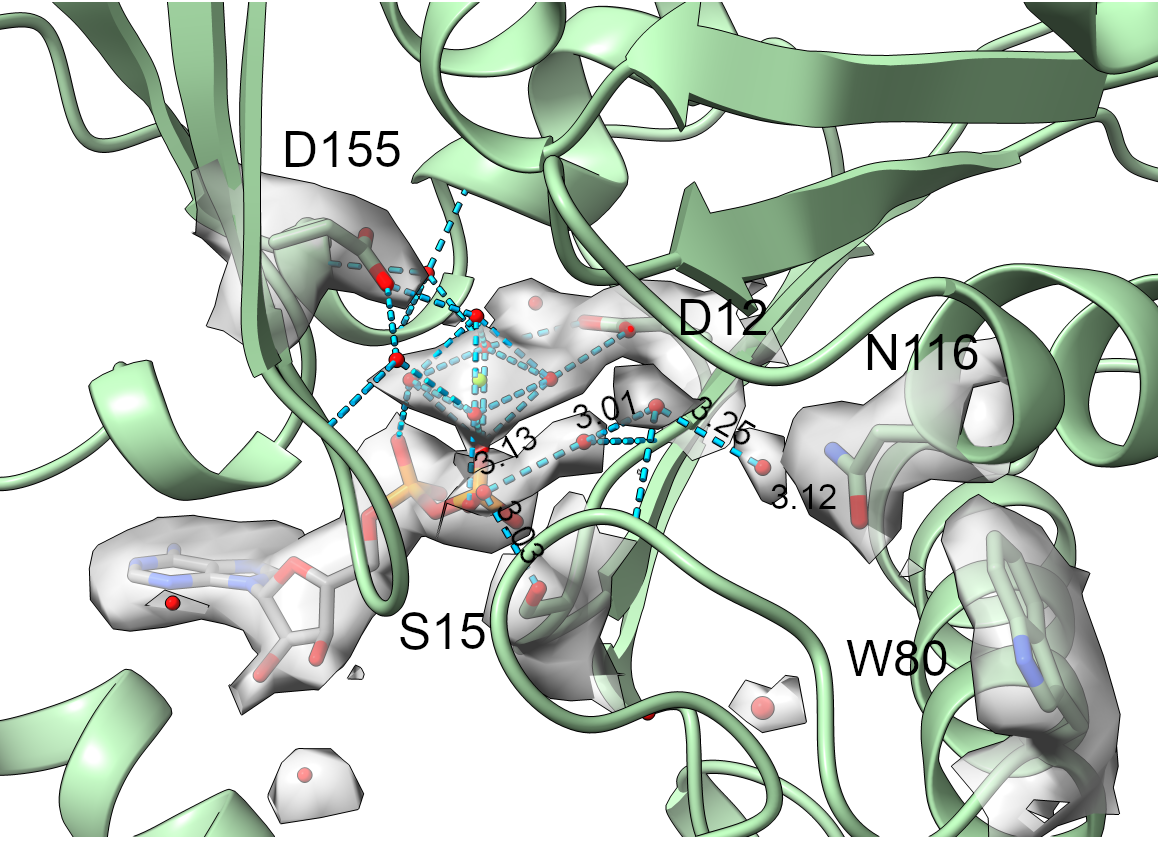

**Figure S7.** Active site in the Act1 filament. The figure shows the density map around the nucleotide-binding site, including ADP (sticks), Mg^2+^ (green sphere) with coordinating water molecules (red spheres), and putative water molecules (red spheres) in the internal cavity.
